# Candesartan treatment preserves learning and working memory in female TgF344-AD rats

**DOI:** 10.1101/2022.06.14.496112

**Authors:** Christopher G Sinon, Kathleen Carter, Jing Ma, Pritha Bagchi, Xiancong Zhang, Peter-Jon C. Williams, Eric B Dammer, Nicholas T Seyfried, Paul S García, Roy L Sutliff, Ihab M Hajjar

## Abstract

**Background:** Targeting the renin angiotensin system, especially with angiotensin receptor II blockers (ARB), and related vascular dysfunction is a promising therapeutic intervention for cognitive impairment including Alzheimer’s Disease (AD). The underlying mechanisms of the effects of ARB is unclear. This study sought to examine if treatment with candesartan, an ARB, affects neurobehavioral manifestation and the underlying neuro- and vascular mechanisms in male and female TgF344-AD rats.

**Methods:** Candesartan or vehicle was administered to TgF344-AD rats (n=127) daily from 12-months to 18-months of age. Behavioral assays (spontaneous alternation test, novel object recognition, water radial arm maze) and neuropathologic assessment were completed along with brain proteome and measures of contractility in 12- and 18-month rat brains.

**Results:** Untreated 18-month TgF344-AD showed impairments in learning and increased perseverative working memory errors on the water radial arm maze (WRAM). These behavioral changes were corrected with candesartan treatment in female rats only. Treatment with candesartan was also associated with improved vascular reactivity and reduced blood pressure in both wild type and TgF344-AD male and female rats. Although there was no effect on amyloid-β, treatment with candesartan reduced whole brain clusterin, an AD-risk associated protein, and GFAP in female TgF344-AD.

**Conclusions:** Our results demonstrate that candesartan administered in the early stages of AD has a sexual dimorphic response in Tgf344-AD rat, where it reduced cognitive disturbances only in female TgF344-AD rat. These effects appear to be independent of changes in blood pressure and amyloid-β reduction and are likely mediated through mechanisms related to clusterin and GFAP pathways.

## Introduction

Alzheimer’s disease (AD), the most common cause of dementia, has an extended preclinical phase lasting decades before clinical phase of the disease[1]. There is currently no known cure for AD and the available treatment options offer only symptomatic relief without altering the progression of the disease[2, 3]. Recent advances using MRI, PET scan, and biochemical analysis of cerebrospinal fluid[4, 5] to identify biomarkers for the initial stages of AD in humans have made it possible to focus on treatment strategies designed to slow the progression or ideally prevent the disease.

The amyloid cascade hypothesis proposes that the cleavage of amyloid precursor protein into amyloid-β, followed by the subsequent aggregation of amyloid-β into destructive extracellular plaques, is the primary cause of AD disease progression[6, 7]. The “vascular hypothesis” of AD is an alternative model suggesting that the development of vascular dysfunction may be an important and early trigger in AD, and potentially a contributor to further amyloid-β aggregation[8, 9]. The evidence for the latter hypothesis has been mixed. One of the primary routes of clearance of amyloid-β from the brain is through the blood brain barrier[10]. However if there is a breakdown in the vasculature of the brain, such as damage to the blood brain barrier, cardiovascular disease, or deposition of amyloid into the blood vessels, this can cause poor clearance of amyloid-β and result in aggregation of amyloid plaques[3]. Targeting the amyloid cascade especially in the symptomatic stages has been mostly disappointing and other targets are being explored.

Therapeutic strategies directed towards the renin angiotensin system (RAS) offer a different, amyloid-β-independent, paradigm for AD therapeutics. Binding of angiotensin II to AT1 receptors causes vasoconstriction, oxidative stress, and endothelial dysfunction [11]. Angiotensin II also increases amyloid-β production by stimulating modulation of APP and increasing γ-secretase activity[12]. Multiple human studies including those conducted by our team suggest that ARBs in general, and candesartan in particular, have neuroprotective effects in early stages of cognitive impairments and AD[13]. However, the underlying mechanisms remain uncertain.

Women are at a higher risk for AD than men and estrogen has been suggested to be protective against both cardiovascular disease and AD[14, 15]. Estrogen is an important regulator of RAS activity, and estrogen deficiency in women and female preclinical models biases RAS activity towards the vasoconstrictive AT1R pathway[16-18]. Estrogen deprivation by bilateral ovariectomy in rats was found to increase AD-related protein expression in hippocampus[19]. A recent study has shown beneficial effects of ARBs to improve cognitive performance and decrease AD biomarkers in an ovariectomized rat model of AD[18].

We undertook a study to determine how chronic ARB treatment starting in the early stages of AD would alter AD progression and vascular dysfunction. These rats display amyloid plaques that precede behavioral impairments, potentially suggesting that it is a preclinical animal model. Additionally, male and female rats were compared for differences in disease progression and response to ARB treatment. The goals of the present work are to assess the effect of candesartan on behavioral, neuropathological, and proteomic mechanisms throughout the early stages of AD in transgenic rats.

## Methods

All experiments were performed at the Atlanta VA Health Care System Animal Facility with approval from the Institutional Animal Care and Use Committee and followed guidelines provided the National Institutes of Health and the Association for the Assessment and Accreditation of Laboratory Animal Care. This manuscript adheres to the applicable Animal Research: Reporting *In Vivo* Experiments (ARRIVE) guidelines[20].

### Animals

Transgenic Fischer 344 rats and wild type littermates were used for data collection. The transgenic Fischer 344 rat model of AD (TgF344-AD rat) was developed using two genes linked to familial Alzheimer’s disease, mutant human *APP*_*SW*_ and presenilin 1 (*PS1ΔE9)* [21]. The rats were housed with a 12-hour light/dark cycles with *ad libitum* access to standard rodent chow and water. Preliminary testing using candesartan in the 3xTgAD mouse revealed a 29% decrease in Aβ1-42 relative to saline (data not shown). With this in mind, a sample size of 10 rats per group provided statistical power to detect an 11% effect of candesartan (power analysis with an α=0.05 and β=0.8). Males and females were analyzed separately to identify sex effects in disease progression and treatment response. A total of 127 rats were bred from an established colony of TgF344-AD rats for data collection. 45 rats were included in the 12-month old cohort and 82 rats were included in the 18-month cohort. Over the course of the study, 7 rats were removed when they met humane endpoints for euthanasia.

### Y-maze Continuous Spontaneous Alternation Test

Rodents introduced to a Y-maze display a tendency to alternate arm entries during maze exploration. This tendency towards alternation is thought to result from a preference towards investigation of new environments, and the spontaneous alternation test takes advantage of this behavior as an assessment of spatial working memory[22]. At 12-months, rats were allowed to freely explore the Y-maze (San Diego Instruments, San Diego, CA, USA) with all three arms open for 8 minutes. The number of arm entries and the path of arm exploration were recorded manually by the experimenter. If at any time the rat remained stationary for >60 seconds during the session, movement was motivated by briefly grasping and releasing the rat from the base of the tail.

A successful alternation was scored when the rat completed three consecutive arm entries via turns in the same direction. Alternation percentage was determined using the following equation[23]:

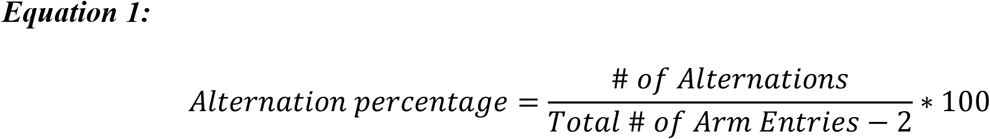

### Novel Object Recognition Test

Following completion of the Y-maze, rats were tested for attention and working memory performance using the novel object recognition test. On the first day, rats were allowed to freely explore a high-walled open field arena for 15 minutes to acclimate to the testing environment. On the second day, rats were again acclimated to the testing environment, followed by a 30-minute break, before starting the training phase of the novel object recognition test. Two identical non-odorous plastic toys (A1 & A2) were affixed with one on each end of the testing environment. Rats were placed into the center of the environment and given seven minutes to explore the testing environment during the training phase. Rats were then returned to their transfer cage for a 20-minute intertrial interval. The choice phase was completed using a third identical plastic toy (A3) and new non-identical plastic toy (B). Each phase was recorded by an overhead video camera and the behavioral data was autoscored using Noldus Ethovision XT11.5 software (Noldus Information Technology, Wageningen, The Netherlands). For each rat, the time spent on each side of the environment and the time spent exploring each object, measured as the amount of time each rat’s nose entered a 2.5-cm radius of the object, was recorded for each session.

The novel object observation time (TN) and the familiar object observation time (TF) were recorded. The discrimination index for the testing phase was calculated by (TN-TF)/(TN+TF). Animals were excluded if they did not cross the midpoint of the arena during any of the phases. Animals were also excluded if they spent more than 80% of total object observation time with either object during the training phase.

### ARB Treatment

From 12- to 18-months of age, all rats on study were treated daily with either the angiotensin receptor II blocker, candesartan, or vehicle (saline). The rats were presented with and without 5 mg/kg of candesartan depending on their experimental groups. Peanut oil proved to be an effective method of daily administration for the ARB treatment. All rats on study consistently consumed all the peanut oil they were offered, regardless of whether it contained candesartan or vehicle. Rats treated with candesartan for one week had no pressor response to an AngII infusion (data not shown).

### Water Radial Arm Maze

The Water Radial Arm Maze (WRAM) consists of eight evenly spaced arms radiating out from a central area. Our protocol for use of the WRAM was adapted from [24]. The maze was filled with opaque water to hide the presence of four escape platforms at the ends of specific maze arms. Unique room cues were placed on the walls around the maze.

At 17.5-months of age, rats were trained to complete the WRAM for 12 consecutive days. Each training session consisted of four trials which were completed when the rat found an escape platform. Before the first session, each rat was randomly assigned escape platform placements that could be navigated using the cues surrounding the maze and which stayed consistent for all 12 sessions.

Upon finding an escape platform, the rat was permitted to remain on the escape platform for 15 seconds and then placed in a cage under a heat lamp for a 30-second intertrial interval. The discovered platform was removed from the maze after each trial. On any trial, if the rat failed to escape the maze within 120-seconds, they were guided to the nearest platform to complete the trial. Therefore, each rat interacted with all four escape platforms during each training session. For each trial, the latency to find an escape platform and the pattern of arm entries during maze exploration was manually scored by the experimenter.

### Blood Pressure Readings

Blood pressure measurements were obtained from the femoral artery at 12-months and 18-months of age in the presence and absence of candesartan treatment. Briefly, rats were anesthetized with isoflurane while maintaining a 37°C body temperature on a thermostatic heating table. The hindlimb region was shaved and disinfected, an incision was made, and the vessel was isolated using blunt dissection, avoiding damaging nerve fibers. The isolated artery was tied distally, and a second suture was looped around the vessel proximal to where the cannula will be inserted. A trochar needle was used to puncture the vessel, and a high-fidelity catheter (Transonic Systems) was inserted into the vessel. Blood pressure was monitored for 15-20 minutes.

### Tissue Collection

Rats were deeply anesthetized using pentobarbital for tissue collection. Arterial blood samples were collected via cardiac stick. Afterwards, cardiac perfusion was performed using ice cold phosphate buffered saline. Rat brains were rapidly extracted and bisected into hemispheres. One hemisphere was fixed in 4% paraformaldehyde and then transferred to a phosphate buffered storage solution containing 0.01% sodium azide. The other hemisphere was snap frozen in liquid nitrogen and stored at −80°C. Aortas were excised and used to study vascular function.

### Vascular Function

Segments of rat aorta, 5 mm in length, from age- and sex-matched littermate controls and TgF344-AD rats, were studied as previously described[25, 26]. Briefly, aortas were isometrically mounted on stainless steel wires, placed in an organ chamber containing Krebs-Henseleit buffer (118 mM NaC1, 4.73 mM KC1, 1.2 mM MgSO_4_, 0.026 mM EDTA, 1.2 mM KH_2_PO_4_, 2.5 mM CaC1_2_, 11 mM glucose, and 25 mM NaHCO_3_; pH when bubbled with 95% O_2_, 5% CO_2_ is 7.4 at 37 °C), and connected to a Harvard apparatus differential capacitor force transducer. For each aorta, tensions were adjusted to 60 mN and then returned to a resting tension 50 mN to approximate an *in vivo* pressure of 100 mm Hg. Data are recorded using PowerLab digital acquisition and analyzed using Chart Software. Results are expressed as mean ± SE. Concentration-response curves were generated to the contractile agents KCl (0-50 mM) and phenylephrine (PE, 0.1nM to 10 µM). Following precontraction with 2-5 µM PE, a concentration which yields 80-90% maximum contraction, relaxation responses will be examined in response to methacholine (1 nM to 100 µM), and the NO donor, sodium nitroprusside (SNP; 1 nM to 100 µM. Relaxations were calculated as a percent of the preconstricted tone induced by phenylephrine. Statistical analyses are performed by two-way analysis of variance repeated measures using Prism software (Graphpad).

### Tissue Sectioning & Immunohistochemistry

Brain tissue sections were embedded in parrafin and sliced at 8 microns. Sections were deparaffinized and immunohistochemically labeled with antibodies to amyloid-β (4G8) and astrocyte activity (GFAP) on a ThermoFisher autostainer. Amyloid-β sections were pretreated with formic acid, blocked with normal serum, and incubated with primary antibody at 1:10,000, then exposed to biotinylated secondary antibody followed by avidin-biotin complex (Vector ABC Elite kit) and developed with diaminobenzidine (DAB). GFAP sections were incubated with primary antibody (anti-GFAP, Dako, Santa Clara, USA) at 1:5000, then exposed to primary antibody enhancer followed by HRP polymer (ThermoScientific UltraVision LP Detection System) and developed with diaminobenzidine (DAB). CP13 sections were pretreated by exposure to hot citrate buffer, blocked with UltraVision protein block, and incubated with primary antibody at 1:2000, then exposed to primary antibody enhancer followed by HRP polymer (ThermoScientific UltraVision LP Detection System) and developed with diaminobenzidine (DAB).

### Neuropathology Analysis

High resolution digital scans of stained brain slices were analyzed using Qupath software[27]. Positive pixel count analysis was performed on scans for brain slices with 4G8 staining for amyloid beta and GFAP staining for astrocyte activity. Positive pixel percentage, a measure of the ratio of positively stained pixels to the total analysis area, was recorded to quantify markers for AD-like pathology in rat brain tissue. Pixels were considered positive for the presence of stain for either amyloid-β or GFAP if the OD unit value was greater than the DAB threshold value (0.4 OD units). Positive cell detection analysis was performed on scans for brain slices with CP13 staining for phosphorylated tau protein (pSer202). Cells were identified as positive for CP13 staining if the cell’s OD mean was above the threshold of 0.4 OD units.

### Sample preparation for mass spectrometry analysis

Frontal cortex samples were collected from 12- and 18-month-old WT and TgF344-AD rats. For 18-month old rats, samples were analyzed from both vehicle and candesartan treated rats. A total of n=4 samples were analyzed for each experimental group. Brain tissues were lysed in 8 M urea lysis buffer (8 M urea, 10mM Tris, 100 mM NaH_2_PO4, pH 8.5) with HALT protease and phosphatase inhibitor cocktail (ThermoFisher). Homogenization was performed using a Bullet Blender (Next Advance) according to the manufacturer’s protocol[28]. Briefly, each tissue piece was added to urea lysis buffer in a 1.5 mL Rino tube (Next Advance) harboring 750 mg stainless steel beads (0.9-2 mm in diameter) and blended twice for 5-minute intervals in the cold room (4 °C). Protein homogenates were transferred to 1.5 mL Eppendorf tubes on ice and were sonicated for 3 cycles of 5 s each at 30% amplitude followed by 15 s on ice. Samples were then centrifuged for 5 min at 12,700 rpm. Protein concentration was determined by bicinchoninic acid (BCA) assay (Pierce) and homogenates (100 μg) were aliquoted. Aliquots were treated with 1 mM dithiothreitol (DTT) and 5 mM iodoacetamide (IAA), each for 30 minutes at room temperature. IAA reaction was carried out in dark. Protein was digested 1:100 (w/w) with lysyl endopeptidase overnight at room temperature. 50 mM ammonium bicarbonate (ABC) was added to the samples to dilute to a concentration of less than 2 M urea, followed by incubation overnight at room temperature with 1:50 trypsin. Peptides were desalted with HLB columns (Waters) and dried under vacuum.

### Liquid chromatography coupled to mass spectrometry

The data acquisition by LC-MS/MS was adapted from a published procedure (Seyfried, Dammer et al. 2017). Derived peptides were resuspended in 100 µL of loading buffer (0.1% trifluoroacetic acid, TFA). Peptide mixtures (2 uL) were separated on a self-packed C18 (1.9 µm, Dr. Maisch, Germany) fused silica column (50 cm x 75 µM internal diameter (ID); New Objective, Woburn, MA) attached to an EASY-nLC™ 1200 system and were monitored on a Q-Exactive HF-X Hybrid Quadrupole-Orbitrap Mass Spectrometer (ThermoFisher Scientific, San Jose, CA). Elution was performed over a 100 min gradient at a rate of 350 nL/min (buffer A: 0.1% formic acid in water, buffer B: 0.1 % formic acid in acetonitrile): The gradient started with 1% buffer B and went to 40% in 100 minutes, then increased from 40% to 99% within 5 minutes and finally staying at 99% for 15 minutes. The mass spectrometer cycle was programmed to collect one full MS scan followed by 20 data dependent MS/MS scans. The MS scans (400-1200 m/z range, 3 × 10^6^ AGC target, 100 ms maximum ion time) were collected at a resolution of 120,000 at m/z 200 in profile mode. The HCD MS/MS spectra (2 m/z isolation width, 30% collision energy, 5 × 10^5^ AGC target, 100 ms maximum ion time) were acquired at a resolution of 15,000 at m/z 200. Dynamic exclusion was set to exclude previously sequenced precursor ions for 20 seconds within a 10 ppm window. Precursor ions with +1, and +8 or higher charge states were excluded from sequencing.

### Label Free Quantification

Label-free quantification analysis was adapted from a published procedure (Seyfried, Dammer et al. 2017). Spectra were searched using the search engine Andromeda, integrated into MaxQuant, against rat Uniprot database (37678 target sequences) plus sequence of mutant human APP_SW_ and PS1ΔE9 proteins. Methionine oxidation (+15.9949 Da), asparagine and glutamine deamidation (+0.9840 Da) and protein N-terminal acetylation (+42.0106 Da) were variable modifications (up to five allowed per peptide); cysteine was assigned a fixed carbamidomethyl modification (+57.0215 Da). Only fully tryptic peptides with up to two miscleavages were considered in the database search. A precursor mass tolerance of ±20 ppm was applied before mass accuracy calibration and ±4.5 ppm after internal MaxQuant calibration. Other search settings included a maximum peptide mass of 6,000 Da, a minimum peptide length of six residues and 0.05-Da tolerance for high resolution MS/MS scans. The FDR for peptide spectral matches, proteins and site decoy fraction was set to 1%. Quantification settings were as follows: requantify with a second peak-finding attempt after protein identification is complete; match full MS1 peaks between runs; use a 0.7-min retention time match window after an alignment function was found with a 20-min retention time search space. The LFQ algorithm in MaxQuant[29, 30] was used for protein quantitation. The quantitation method considered only razor and unique peptides for protein level quantitation.

Two samples from the cohort were identified as outliers via multidimensional scaling and were removed from further analyses (one sample each from the 12-month/Male/WT group and the 18-month/Female/WT/Vehicle group). The log_2_ LFQ abundance values from these samples were highly dissimilar from all other samples. We implemented a median polish algorithm for removing technical variance (for example, due to tissue collection, cohort or batch effects) from a two-way abundance-sample data table as originally described by Tukey[31]. Detailed methods have been published previously[32] and the algorithm is fully documented and available as an R function that can be downloaded from https://github.com/edammer/TAMPOR.

From a final list of 36655 peptides belonging to 3801 protein groups, 3301 proteins were identified where fewer than 50% of LFQ intensity measurements were missing within every experimental group. Missing values were resolved using Perseus-style imputation method based on the assumption that log_2_ LFQ abundances would be normally distributed. Imputed values were selected from a normal distribution (with +/- 0.3 SD) centered at −1.8 SD from the mean LFQ abundance.

### Statistical Analysis

Statistical testing was performed using Prism 7.0 (GraphPad Software, San Diego, CA, USA) and IBM SPSS Statistics V26 (IBM Corporation, Armonk, NY, USA). A sample size of 10 rats per group was justified based on preliminary data collected in the 3xTgAD mouse model. Preliminary data in this model showed a 29% decline in amyloid-β accumulation. Power analysis in G*Power 3.1.9.4 with α=0.05, β=0.80 and an anticipated effect size of 0.8 was adequate to detect a minimal effect of candesartan of 11%. All datasets were tested for normality using the Shapiro-Wilk test. Tests for normality perform poorly with sample sizes comparable to those used in these experiments, so we compared the Shapiro-Wilk test results against visual inspection of the data distributions in the selection of non-parametric vs. parametric tests. Neuropathology data were analyzed with Student’s t-test with Welch’s correction or Two-way ANOVA with Sidak’s test for multiple comparisons. One-way ANOVA was used to analyze changes in neuropathology between 12- and 18-months of age. Student’s t-test for independent samples was used for analysis of data from the continuous spontaneous alternation test. For the NOR, the 95% confidence interval was computed for each group. We determined that each group recognized the novel object if the 95% confidence interval did not cross the line at DI=0.0 (signifying performance at the level of chance). MAP data was analyzed by one-way ANOVA. WRAM data was analyzed using two-way ANOVA with Sidak’s test for multiple comparisons. Planned contrasts to analyze drug effects on learning and memory performance were analyzed using Sidak’s test for multiple comparisons. Differential protein expression was determined by one-way ANOVA followed by Tukey’s comparison post-hoc test between all experimental groups. Volcano plots were used to display significant alterations in protein expression between groups. The log-transformed fold change (Log_2_ Difference) between groups was plotted on the x-axis, and the Tukey adjusted ANOVA *p*-value (-Log_10_ *p*-value) was plotted on the y-axis (α=0.05).

## Results

### No differences in continuous spontaneous alternation test performance or novel object recognition at 12-months in TgF344-AD or WT rats

There were no observed differences in spatial working memory between 12-month-old WT or TgF344-AD rats on the continuous spontaneous alternation test. Spontaneous alternation percentage within the Y-maze did not differ between strains (Student’s t-test, p=0.908, ns, Figure 1A). There was also no difference seen in the number of arm entries observed during maze exploration (Student’s t-test, p=0.681, ns, Figure 1B).

**Figure 1.**
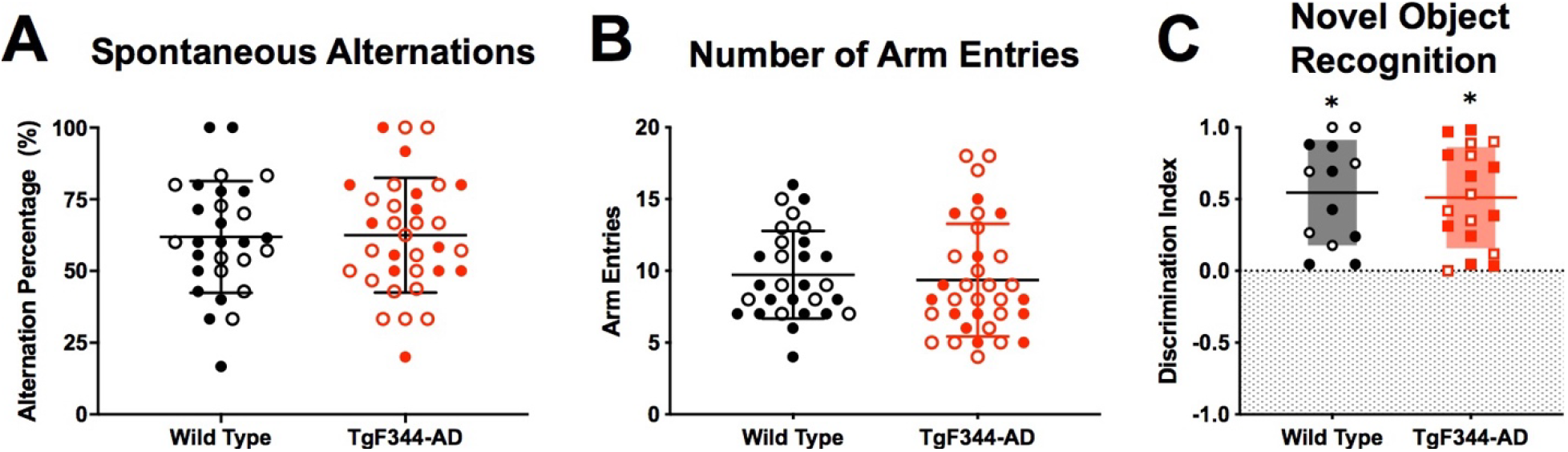
No difference in spontaneous alternation test performance between TgF344-AD rats and wild-type littermates at age 12-months. A) There was no difference in alternation behavior observed between TgF344-AD (n = 34) and wild-type (n = 29) rats (Student’s t-test, p=0.908). B) There was no difference in the number of arm entries made during maze exploration between TgF344-AD (n = 34) and wild-type (n = 29) rats (Student’s t-test, p=0.681). Open symbols indicate female rats and closed symbols indicate male rats. C) Both TgF344-AD (N=9) and wild-type rats (N=10) can discriminate the novel object from a familiar object during the testing phase of the novel object recognition test. Results are shown as a mean with the shaded region representing the 95% confidence interval. The 95% confidence interval for the discrimination index of both wild-type (DI range = 0.119-0.650) and TgF344-AD (DI range = 0.057-0.693) rats are above 0. Therefore performance for both groups was better than chance demonstrating preference for the novel object. Open symbols indicate female rats and closed symbols indicate male rats.

Both WT and TgF344-AD rats were able to recognize the novel object during testing sessions performed at 12-months of age. Results are presented as the calculated group mean for the discrimination index, with error bars representing the 95% confidence interval of the mean. A discrimination index equal to 0 represents performance at chance for recognition of the novel object. Both wild-type (DI range = 0.119-0.650) and TgF344-AD (DI range = 0.057-0.693) rats were able to discriminate the novel from the familiar object during the testing phase (Figure 1C). Performance did not differ between males and females on either test, so both sexes were grouped together for analysis.

### Candesartan improves learning on the WRAM for 18-month female, but not male, TgF344-AD rats

The WRAM can be used to assess learning, as well as working and reference memory, since performance on the task will ideally improve from session 1 to session 12. To assess learning on the WRAM in WT and TgF344-AD rats, the cumulative latency to reach the escape platforms and the total combined errors of the initial and final training sessions were compared. Data from sessions 1 & 2 were combined for the “initial” training data. Likewise, data from sessions 11 & 12 were combined for the “final” training data.

Both WT and TgF344-AD rats reduced their time to complete the maze over the course of 12 training sessions. All groups reduced their time to complete the maze, regardless of prior treatment with vehicle or candesartan. For male and female WT rats, there was a significant reduction in the latency to reach all escape platforms as measured by two-way ANOVA with repeated measures (main effect of time, F(1,15)=96.67,p<0.001 & F(1,14)=41.33,p<0.001, Figure 2A&B respectively). Planned contrasts using Sidak’s multiple comparison test revealed a significant improvement in time to complete the maze on the final sessions for WT rats receiving vehicle (Males - Initial: M=470 sec, Final: M=142 sec; Females - Initial: M=355 sec, Final: M=124 sec) and candesartan (Males - Initial: M=464 sec, Final: M=90 sec; Females - Initial: M=477 sec, Final: M=140 sec). There was a similar reduction in latency to reach all platforms across training sessions for male and female TgF344-AD rats (main effect of time, F(1,12)=32.54,p<0.001 & F(1,19)=29.2,p<0.001, Figures 2E&F respectively). Post-hoc analysis showed that TgF344-AD rats receiving vehicle (Males - Initial: M=480.6 sec, Final: M=239.8 sec; Females – Initial: M=341.5 sec, Final: M=194.9) and candesartan (Males - Initial: M=437.7, Final: M=198.8; Females – Initial: M=396.8, Final: M=152.6) completed the maze more quickly over the course of training.

**Figure 2.**
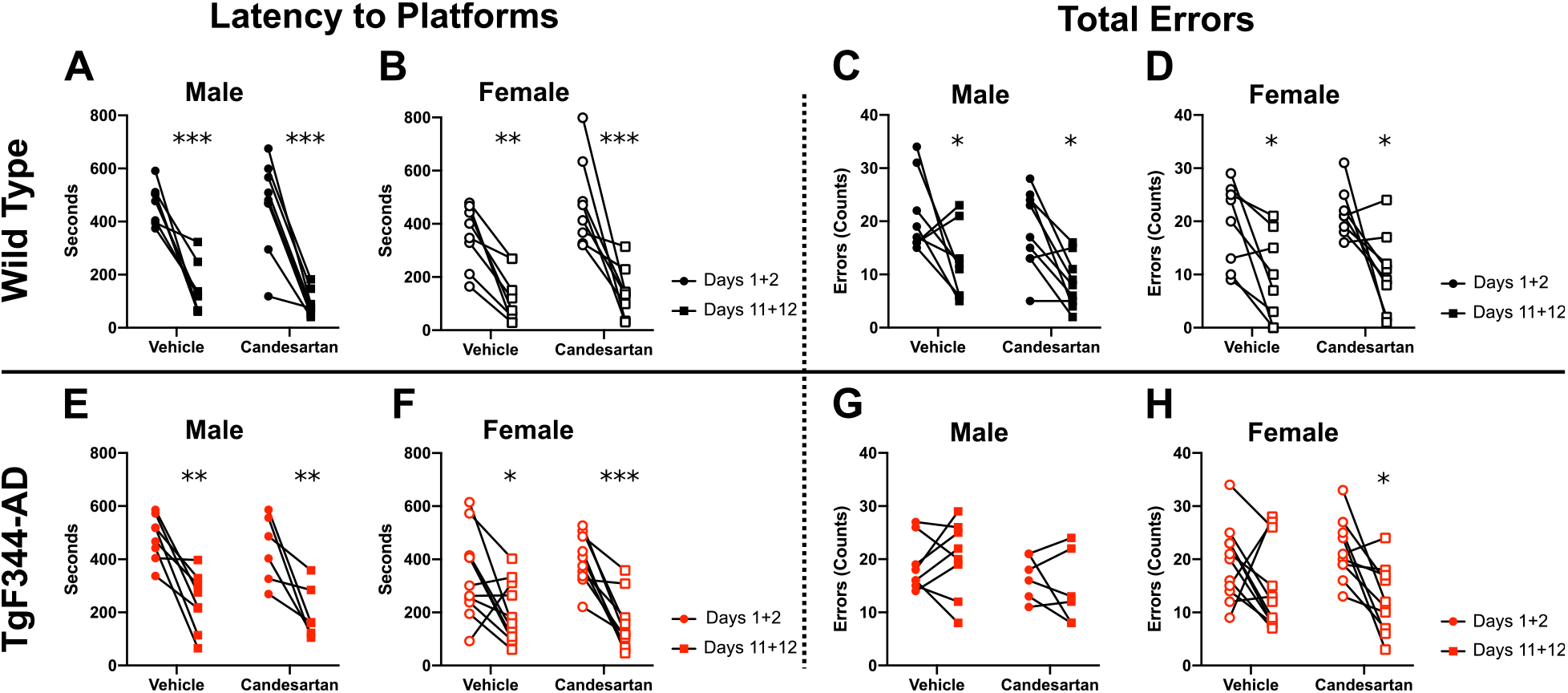
Candesartan improves WRAM maze performance in female TgF344-AD rats. A & B) Repeated measures comparisons revealed that there was a significant decrease in the time male and female WT rats took to complete the WRAM over the course of 12 days of training (Sidak Multiple Comparisons Test, [A] ***,p<0.001 & ***,p<0.001; [B] **,p<0.005 & ***,p<0.001). C & D) Male and Female WT rats also learned to complete the WRAM with fewer total errors (Sidak Multiple Comparisons Test, [C] *,p=0.036 & *,p=0.019; [D] *,p=0.026 & *,p=0.015). E & F) Male and femaleTgF344-AD rats also decrease the time needed to complete the WRAM by the final sessions (Sidak Multiple Comparisons Test, [E] **,p=0.002 & **,p=0.006; [F] *,p=0.017 & ***,p<0.001). G & H) However, only female TgF344-AD rats treated with candesartan learned to decrease their total cumulative errors by the final sessions. TgF344-AD rats that were administered vehicle committed as many total errors on their final sessions as they had in the initial training sessions ([G] Repeated Measures Two-way ANOVA, F(1,12)=0.067,p=0.800; [H] Sidak Multiple Comparisons Test, vehicle: ns,p=0.260, candesartan: *,p=0.017).

WT rats learned to complete the WRAM with fewer cumulative working memory errors, reference memory errors and perseverative errors between the initial and final sessions in the WRAM (main effect of time, F(1,15)=15.79,p=0.001 & F(1,14)=17.89,p=0.001, Figure 2C&D). There was a significant reduction in errors over the course of training for groups receiving vehicle (Male - Initial: M=21.25, Final: M=12.13; Female – Initial: M = 19.50, Final: M = 9.375) and candesartan (Male - Initial: M=18.11, Final: M=8.44; Female – Initial: M = 21.63, Final: M = 10.50) as determined by planned contrasts using Sidak’s multiple comparison test.

However, the response to candesartan in TgF344-AD rats was sex dependent. Male rats did not reduce their total maze errors after training (no main effect of time, F(1,12)=0.067,p=.8002, Figure 2G). Planned contrasts found no difference in the errors accumulated for male TgF344-AD rats treated with vehicle (Initial: M = 18.88, Final: M = 20.13) or candesartan (Initial: M = 16.67, Final: M = 14.50). Though male TgF344-AD rats were unable to learn to complete the maze with fewer errors, female TgF344-AD improved their maze performance after treatment with candesartan. There was an overall reduction in errors after training for all TgF344-AD rats (Main effect of time, F(1,19)=10.21,p=0.005). This effect was driven by significant reduction in total errors from rats treated with candesartan (Initial: M=21.70, Final: M=12.40). Female TgF344-AD rats receiving vehicle showed no reduction in the number of errors performed on the WRAM following training (Initial: M=19.27, Final: M=14.64, Figure 2H).

### Amyloid-β and GFAP, but not p-Tau, are increased in the hippocampus of 12-month old TgF344-AD rats with no associated behavioral changes

Coronal slices taken from approximately −2.5mm posterior from bregma were used to quantify amyloid-β and GFAP staining at the level of the dorsal hippocampus (Figure 3A-B,G-H). At 12-months old, amyloid-β accumulation was already apparent in the brains of TgF344-AD rats. Image analysis revealed that there were significantly more pixels that were positive for amyloid-β staining in the hemisphere slices taken from TgF344-AD rats than from WT rats (Student’s t-test with Welch’s correction, p<0.001, Figure 3C). Additionally, there was an increase in GFAP staining across the brain slices collected from TgF344-AD rats when compared with WT rats (Student’s t-test with Welch’s correction, p<0.013, Figure 3I). Similar results were found when quantification was focused on the region of dorsal hippocampus specifically. Both amyloid-β (p<0.001, Figure 1E) and GFAP (p=0.004, Figure 3K) staining was increased in the dorsal hippocampus of TgF344-AD rats. Phosphorylated tau protein levels were unchanged in TgF344-AD rats at 12-months. There was no significant differences in the percentage of positively stained cells within the hippocampal pyramidal cell layer between WT and TgF344-AD rats. The percentage of stained cells was similar in both CA1 (Figure 3O, Student’s t-test with Welch’s correction, p=0.532) and CA3 (Figure 3P, Student’s t-test with Welch’s correction, p=0.806).

**Figure 3.**
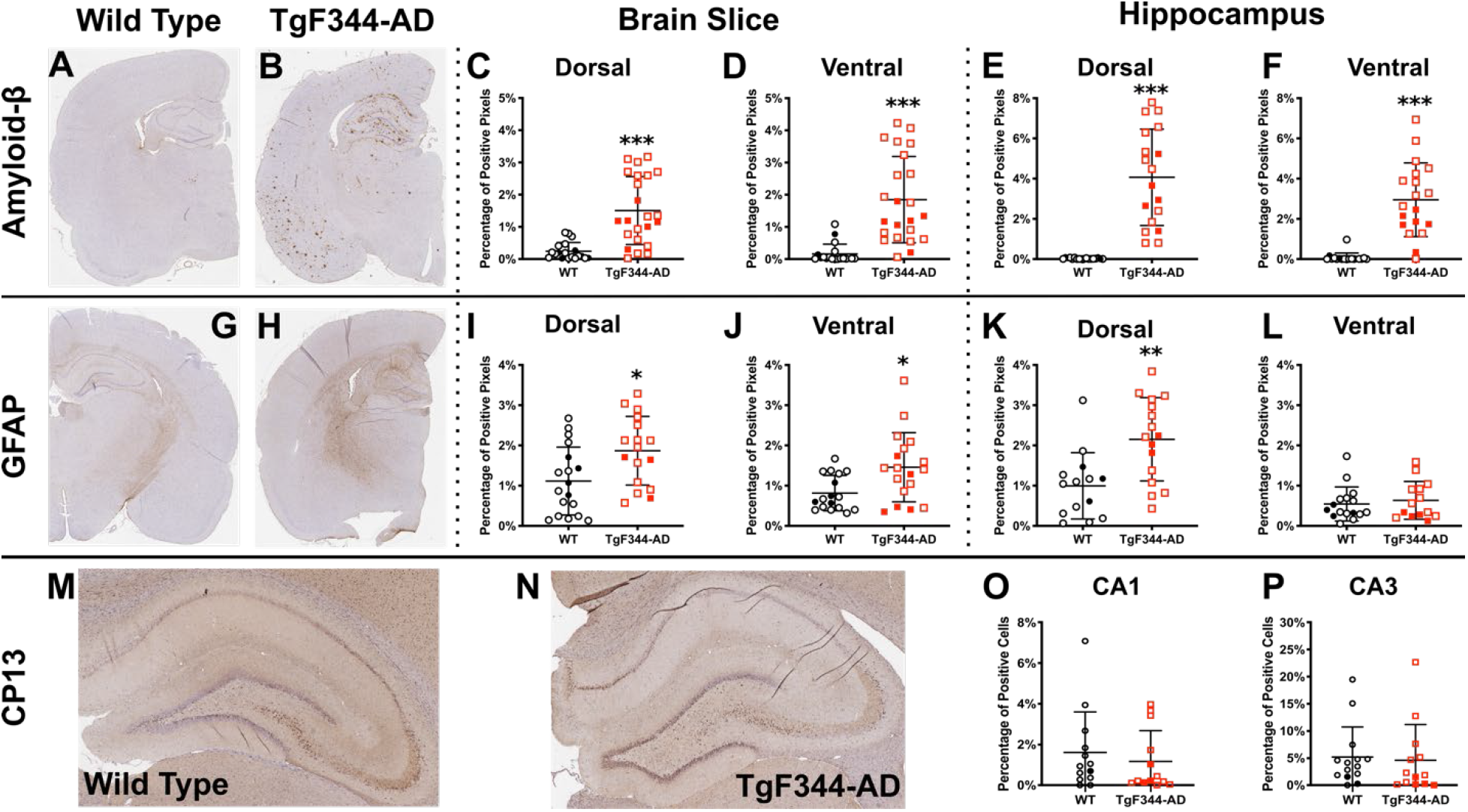
Amyloid-β and GFAP levels are increased in TgF344-AD rats at age 12-months. A-B) Examples of stained brain slices used for quantification of amyloid-β. DAB staining for amyloid-β and GFAP appears brown on the slides. C-F) TgF344-AD rats display increased accumulation of amyloid-β throughout the entire slide and localized specifically to hippocampus at 12-months (Student’s t-test with Welch’s correction, p<0.001 for all). G-H) Examples of stained brain slices used for quantification of GFAP. I-J) There was a general increase in GFAP throughout the brains of TgF344-AD rats at 12 rats. K) However, only the dorsal hippocampus displayed an increase in GFAP over the WT (Student’s t-test with Welch’s correction, p=0.004). L) There was no difference seen between strains in the amount of GFAP present in ventral hippocampus. M-N) Examples of stained dorsal hippocampus sections used for quantification of phosphorylated tau protein. O-P) There were no differences seen between strains in the amount of stained phosphorylated tau protein (CP13) within cells identified in the CA1 or CA3 pyramidal cell layer. Open symbols indicate female rats and closed symbols indicate male rats.

Quantification of amyloid-β and GFAP was also completed in coronal slices made at the level of ventral hippocampus, approximately −5.0mm posterior from bregma. Amyloid-β was found to be significantly increased in TgF344-AD rats across the slice (p<0.001, Figure 3D) and within ventral hippocampus specifically (p<0.001, Figure 3F). However, while GFAP was increased for TgF344-AD rats across the slice (p=0.012, Figure 3J), there was no differences in staining for GFAP within the ventral hippocampus of WT and TgF344-AD rats (p=0.798, Figure 3L).

### Brain β-amyloid and GFAP levels increased in 18-month old TgF344-AD rats

Coronal sections were taken from WT and TgF344-AD rats at 18-months (Figure 4A-B). Amyloid-β was significantly increased in the dorsal hippocampus of TgF344-AD when compared with WT rats (main effects of genotype, [F(1,46)=140.4,p<0.001, Figure 4C]). The same relationship is observed in stains for amyloid-β in coronal slices taken at the level of ventral hippocampus (Figure 4D). Additionally, amyloid-β was found to be increased throughout the entire hemisphere at both slice locations compared with WT rats [F(1,60)=167.4,p<0.001, Figure 4G] and [F(1,61)=182.7,p<0.001,Figure 4H] respectively). Post-hoc analysis found all comparisons between WT and TgF344-AD rats to be significant (p<0.001, for all), regardless of the drug treatment for any group. Overall at 18-months of age, there is a negligible signal for amyloid-β in slices from the WT rats and a significant presence of staining for amyloid-β in slices from the TgF344-AD rats.

**Figure 4.**
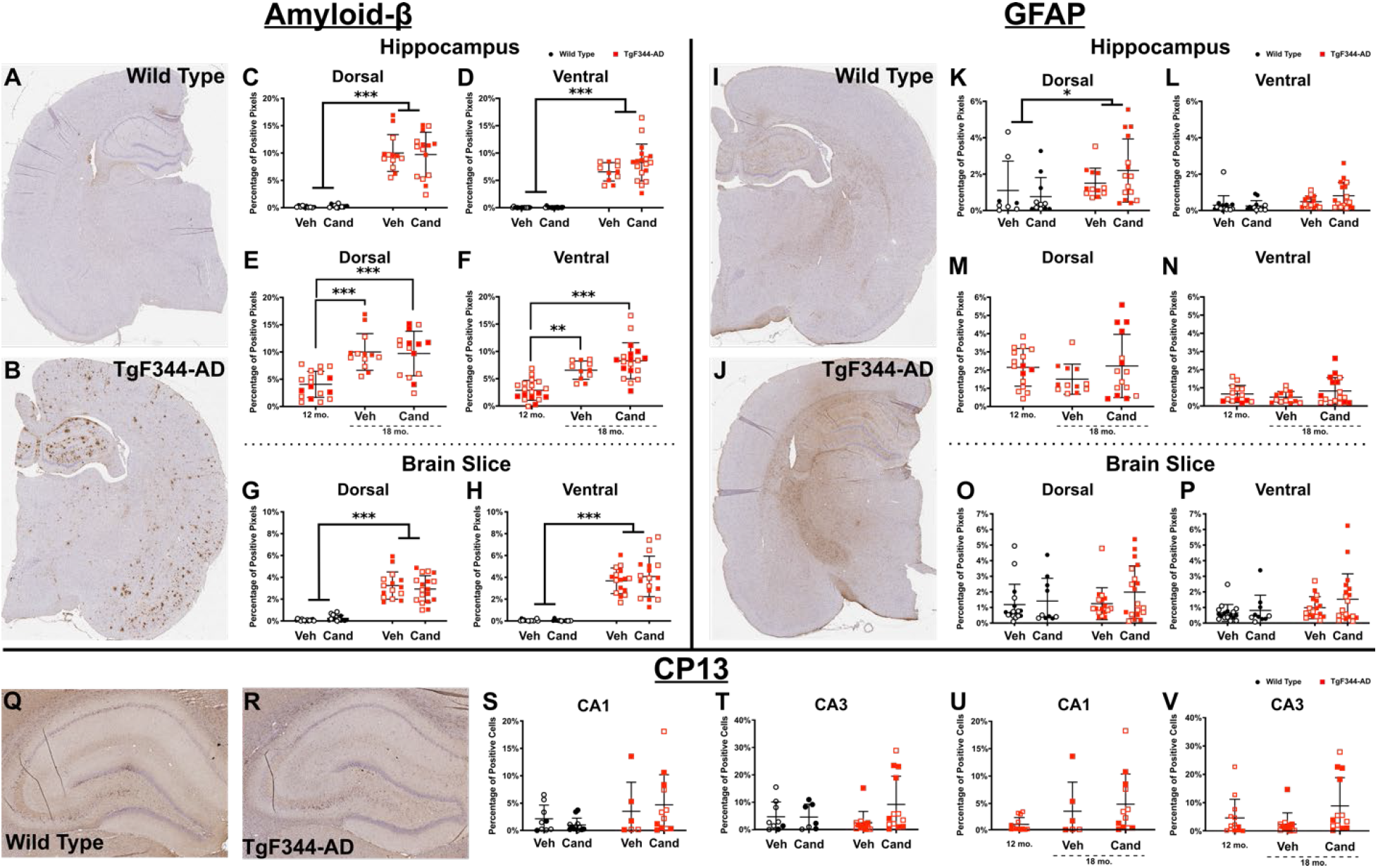
Treatment with candesartan did not affect the accumulation of amyloid-β and GFAP in 18 month old TgF344-AD rats. A-B) Examples of stained brain slices used for quantification of amyloid-β at 18-months. C-D) Two-way ANOVA revealed a main effect of genotype for all comparisons of amyloid-β accumulation in WT and TgF344-AD rats at 18-months. Post-hoc tests revealed significant differences between all WT and TgF344-AD experimental groups (not pictured). E) At 18-months there is a significant increase in amyloid-β in the dorsal hippocampus of TgF344-AD rats compared with levels seen at 12-months (Kruskal Wallis test, p<.001). Multiple comparisons confirmed that this increase is seen in both rats receiving vehicle and those receiving candesartan. F) Amyloid-β is also increased in the ventral hippocampus of TgF344-AD rats when comparing the 12- and 18-month cohorts. G-H) Amyloid-β was increased across the whole hemisphere at both dorsal and ventral locations in TgF344-AD rats. K-L) There was a significant main effect of genotype on the amount of GFAP staining in the dorsal hippocampus of 18-month old rats. Similar to the results at 12-months, there is a significant increase in GFAP in the dorsal hippocampus of TgF344-AD rats relative to WT rats, but there was no observed differences in GFAP within the ventral hippocampus of TgF344-AD and WT rats. M) There were no significant differences in the amount of GFAP seen in the dorsal hippocampus of TgF344-AD between any groups at 12- or 18-months of age. N) There were also no differences in the amount of GFAP noticed in the ventral hippocampus due to time or drug treatment. I-J) There were no significant differences between WT and TgF344-AD in total hemisphere GFAP at either sectioning location. Q-R) Examples of stained dorsal hippocampus sections used for quantification of phosphorylated tau protein. S-T) There were no significant differences between strains in the percentage of stained cells in either the CA1 or CA3 pyramidal cell layer. U-V) There was no difference in the phosphorylated tau staining in either CA1 or CA3 between 12-month and 18-month (treated or untreated) TgF344-AD rats. Open symbols indicate female rats and closed symbols indicate male rats.

Brain slices stained for GFAP at 18-months are presented in Figures 4I-J. For TgF344-AD rats, there was an increase in GFAP found within the region of dorsal hippocampus (main effect of genotype, F(1,42)=4.862,p=0.033, Figure 4K). However, there were no observed differences in GFAP quantification within ventral hippocampus (Figure 4L). There were also no observed differences in GFAP between WT and TgF344-AD rats across the full brain slice at the level of the dorsal or ventral hippocampus (Figure 4O-P).

Representative images of dorsal hippocampus used for CP13 stain detection are shown in Figure 4Q-R. There were no differences in phosphorylated tau found due to strain or ARB treatment in the CA1 or CA3 pyramidal cell layer (Figures S & T, p>0.05 for main effects and interaction effects in CA1 & CA3).

Brain slices from rats sacrificed at 12-months and 18-months were analyzed to determine whether there were changes in the relative amounts of amyloid-β and GFAP present in the brain at different timepoints of the disease in TgF344-AD rats. There were significant differences in amyloid-β staining observed between 12-month TgF344-AD rats, 18-month TgF344-AD rats receiving vehicle, and 18-month TgF344-AD rats receiving candesartan as determined by a Kruskal-Wallis test (p<0.001). Multiple comparisons revealed there was more diffuse amyloid-β staining in both 18-month experiment groups when compared with the 12-month rats. There were no observed differences in stain quantification between the two 18-month groups that could be explained by treatment with candesartan. Similar results were observed at the level of dorsal and ventral hippocampus (Figure 4E-F). The percentage of pixels positively stained for GFAP did not change in the whole slice during disease progression (Figure 4M-N). There was also no change in the percentage of cells stained positive for CP13 in CA1 (p=0.090) or CA3 (p=0.298) between 12-month and 18-month TgF344-AD rats (Figure(U-V).

Tau protein expression was also assayed by comparing the expression of two MAPT isoforms between WT and TgF344-AD experimental groups. Identified peptides were quantified using peptide ion intensities. Using label-free quantification, we were able to quantify a total of 36655 peptides belonging to 3801 protein groups. In total, our analysis mapped 3301 unique gene symbols across a total of 46 samples. Two samples were removed by outlier analysis due to highly dissimilar log_2_ LFQ abundances when compared with the rest of the samples.

Tau protein expression was compared for 12- and 18-month old WT and TgF344-AD rats. Two isoforms of MAPT were identified via label free quantification following liquid chromatography mass spectrometry (LC-MS/MS). MAPT|D3ZKD9 and MAPT|D4A1Q2 expression levels were compared between all experimental groups (Table 1). The lowest *p*-value for any ANOVA-Tukey pairwise comparison was 0.52, so MAPT was not found to be significantly different between WT and TgF344-AD rats in our analysis.

**Table 1.** Summary of tau protein expression in WT & TgF344-AD rats.

|  | MAPT\|D3ZKD9 |  | MAPT\|D4A1Q2 |  |
| --- | --- | --- | --- | --- |
| ANOVA Results |  |  |  |  |
| F | 0.527 |  | 0.837 |  |
| Pr(<F) | 0.871, *n.s.* |  | 0.606, *n.s.* |  |
| Planned Contrasts | Log<sub>2</sub> Difference | *p*-value | Log<sub>2</sub> Difference | *p*-value |
| *Male* |  |  |  |  |
| 12 mo. WT vs. 12 mo. TgF344-AD | -0.549 | 0.992, *n.s.* | -0.087 | 0.999, *n.s.* |
| 12 mo. TgF344-AD vs. 18 mo. TgF344-AD+Veh | -0.277 | 0.999, *n.s.* | -0.228 | 0.847, *n.s.* |
| 18 mo. WT+Veh vs. 18 mo. TgF344-AD+Veh | -0.057 | 0.999, *n.s.* | 0.085 | 0.999, *n.s.* |
| 18 mo. WT+Veh vs. 18 mo. TgF344-AD+Cand | 0.041 | 0.999, *n.s.* | 0.051 | 0.999, *n.s.* |
| 18 mo. TgF344-AD+Veh vs. 18 mo. TgF344-AD+Cand | 0.098 | 0.999, *n.s.* | -0.035 | 0.999, *n.s.* |
| *Female* |  |  |  |  |
| 12 mo. WT vs. 12 mo. TgF344-AD | 0.283 | 0.999, *n.s.* | 0.097 | 0.999, *n.s.* |
| 12 mo. TgF344-AD vs. 18 mo. TgF344-AD+Veh | 0.083 | 0.999, *n.s.* | -0.132 | 0.997, *n.s.* |
| 18 mo. WT+Veh vs. 18 mo. TgF344-AD+Veh | -0.291 | 0.999, *n.s.* | 0.208 | 0.945, *n.s.* |
| 18 mo. WT+Veh vs. 18 mo. TgF344-AD+Cand | 0.239 | 0.999, *n.s.* | 0.100 | 0.999, *n.s.* |
| 18 mo. TgF344-AD+Veh vs. 18 mo. TgF344-AD+Cand | 0.531 | 0.989, *n.s.* | -0.108 | 0.999, *n.s.* |

### Candesartan improved vascular reactivity and decreased mean arterial pressure for both WT and TgF344-AD rats

Vascular reactivity was assessed in thoracic aortic rings from WT and TgF344-AD rats at 12-months and 18-months of age. Vascular contractility was assessed in aortic rings by measuring isometric force in response to increasing concentrations of the depolarizing agent, KCl (10-80mM), and α adrenoceptor agonist, phenylephrine (1nM-30µM). Relaxation was also measured in thoracic aortas after preexposure to 0.3µM phenylephrine. Percentage relaxation was determined following exposure to methacholine (100pM-10µM) and sodium nitroprusside (SNP, 100pM-300nM).

At 12-months, there was no significant difference in vascular contractility in response to KCl between WT and TgF344-AD rats (KCl EC50 [M±SEM]: WT = 29.87±0.39mM, TgF344-AD = 30.49±0.35mM, p=0.250). The EC50 for aortic ring contraction to phenylephrine was increased in TgF344-AD rats (PE EC50: WT = 211±10.0nM, TgF344-AD = 240±9.14nM, p=0.046,*). Vessels from TgF344-AD rats required lower concentrations of methacholine and SNP for relaxation (Methacholine EC50: WT = 277±25.72nM, TgF344-AD = 205±19.45nM, p=0.036,*; SNP EC50: WT = 28.7±2.26nM, TgF344-AD = 20.3±1.88nM, p=0.009,**).

Treatment with candesartan led to similar effects on vascular reactivity for WT and TgF344-AD rats. At 18-months, there were no significant interaction effects between genotype and ARB treatment on vascular reactivity (p>0.05 for all, two-way ANOVA). Planned contrasts between WT and TgF344-AD rats with Sidak’s multiple comparison test found no change in the contractility or relaxation response to any stimulus after six months of treatment with either vehicle or candesartan (adjusted familywise p>0.05 for all). Aortic rings from TgF344-AD rats demonstrated reduced contractility in response to KCl (main effect of genotype, p=0.040,*), and contractility increased in response to phenylephrine (main effect of genotype, p=0.024,*). The main effect of ARB treatment was significant in all vascular reactivity responses (p<0.05 for all). Vessel contraction was stimulated by lower concentrations of KCl and phenylephrine in rats treated with candesartan, however stimulation of the relaxation response required higher concentrations of methacholine and SNP.

Following six months of daily treatment with candesartan or vehicle, there was a main effect of drug treatment on median arterial pressure (One-way ANOVA, p<0.01, Figure 5C). Rats receiving regular doses of an ARB displayed a decrease in the recorded mean arterial pressure when compared with the rats receiving vehicle. The effect of candesartan to decrease mean arterial pressure was not different between WT and TgF344-AD rats.

**Figure 5.**
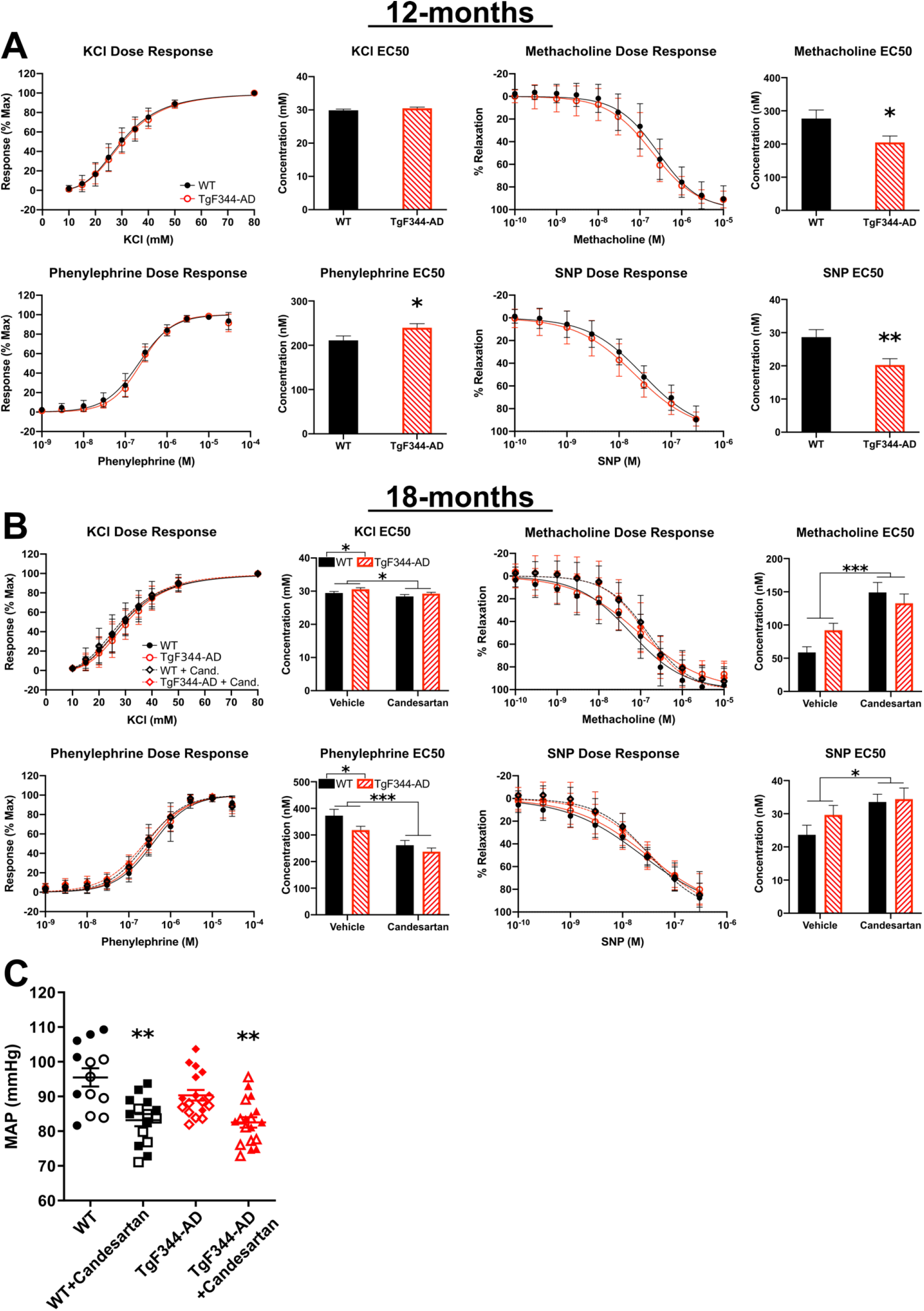
Candesartan alters vascular reactivity and decreases mean arterial pressure for both WT and TgF344-AD rats. A) WT and TgF344-AD rats show slight differences in vascular reactivity at 12-months of age, highlighted by reduced contractility in response to phenylephrine and increased relaxation responses to methacholine and SNP in TgF344-AD rats (p<0.05 for all). B) There was a consistent effect of candesartan to improve vascular reactivity for rats at 18-months (significant main effect of treatment, two-way ANOVA, p<0.05). In WT and TgF344-AD rats treated with candesartan, vascular contractility was reduced in TgF344-AD following response to the depolarizing agent, KCl, and contractility was increased for the α adrenoceptor agonist, phenylephrine. C) Following six months of daily treatment with candesartan or vehicle, there was a main effect of drug treatment on mean arterial pressure. Candesartan effects on mean arterial pressure were not different between WT and TgF344-AD rats. There was a significant difference between treated and untreated groups for each strain (p<0.01, **) as determined by one-way ANOVA. Open symbols indicate female rats and closed symbols indicate male rats, n=13-18. All data expressed as mean±SEM.

### Brain proteome of 18-month TgF344-AD and the effect of candesartan treatment

Protein expression was compared across all groups of WT and TgF344-AD rats at age 18-months. The average protein expression levels for all experimental groups were analyzed by ANOVA with Tukey’s multiple comparisons test for pairwise comparisons. A total of 161 proteins were found to be significantly altered between all groups via one-way ANOVA (*p*<0.05, expression levels for selected proteins displayed in Figure 6). TgF344-AD rats express human APP with Swedish mutation, so it was expected that human APP levels were significantly increased compared with WT rats (F = 36.49, *p* < 0.001). Comparisons between groups also routinely found altered expression of APOE and Midkine (MDK) in TgF344-AD rats. Increased levels of these proteins are associated with an increased risk for AD in humans[33]. Additionally, there were sex differences in GFAP expression in the TgF344-AD rats. One-way ANOVA determined that there was an increase in GFAP expression for male TgF344-AD rats after treatment with candesartan (*p*=0.046), however female TgF344-AD rats did not display significantly more GFAP than WT females following candesartan treatment (*p*=0.240). Our results are summarized below in Table 2, with the relative levels of ß-Actin included for reference.

**Figure 6.**
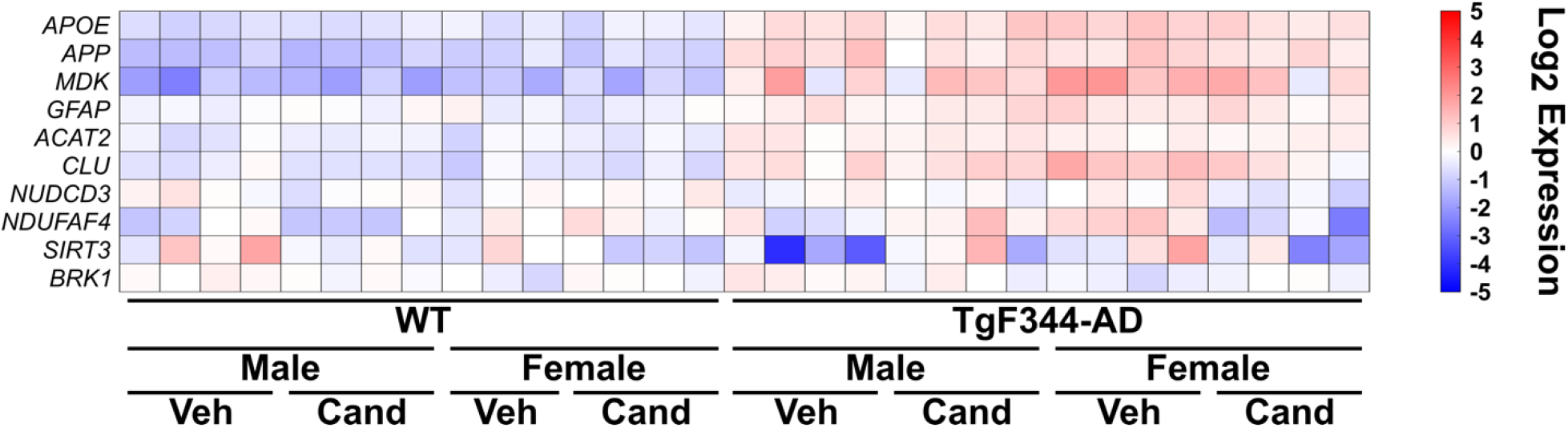
Expression of 161 proteins were significantly altered in 18-month old TgF344-AD rats. Heatmap showing the relative abundance of 10 proteins that were significantly altered between group comparisons.

**Table 2.**
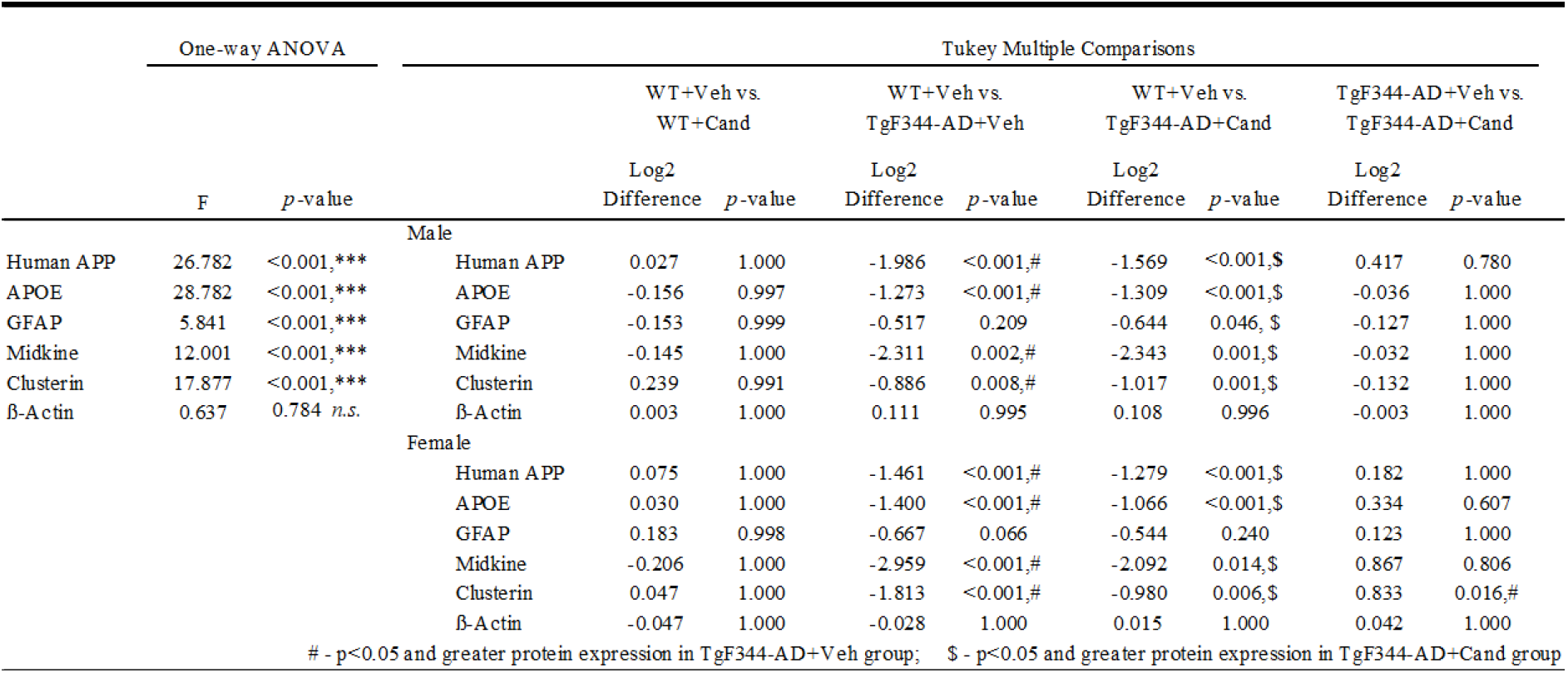
AD-associated protein expression in 18-month WT & TgF344-AD rats.

The effects of candesartan on protein expression were tested in WT rats brains (Figure 7A&B). Among males only one protein displayed a significant difference in expression. A slight increase in expression of V1G1 (ATP6V1G1), a subunit of V-ATPase[34], was found in males following six months of treatment with candesartan. In females, a total of 10 proteins displayed differential expression between vehicle and candesartan treated groups. Following candesartan treatment, there is increased expression of genes associated with protein translation and elongation, such as CAPRIN1 and EEF1B2. Rats treated with vehicle have comparatively more expression of AKAP12, a regulator of PKA activity in endothelial cells, and FAM210A, involved in the regulation of muscle and bone strength.

**Figure 7.**
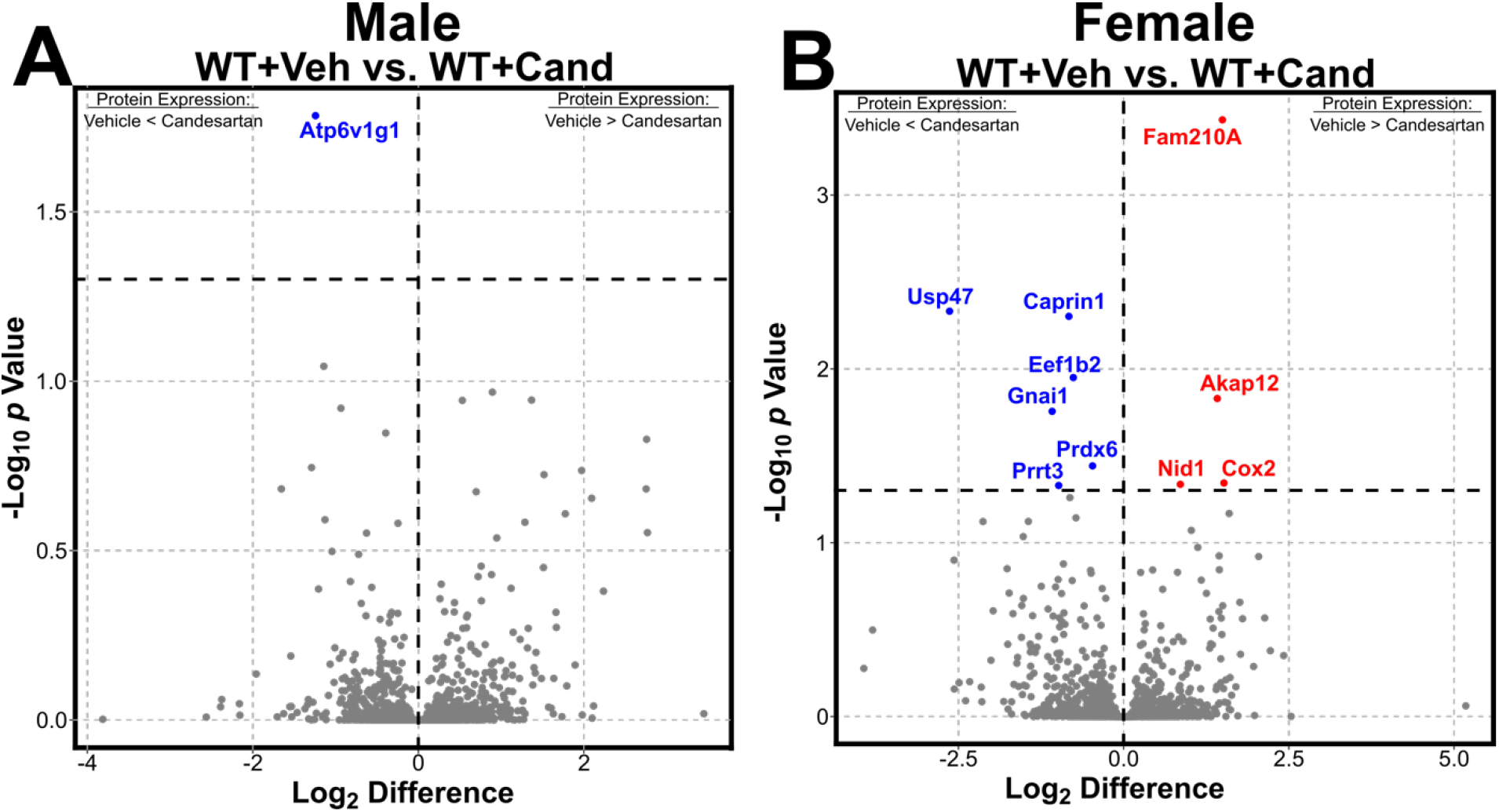
Significantly altered proteins due to candesartan in WT male and female rats. A) One protein, ATP6V1G1, is increased in wild type male rats following six months of daily treatment with candesartan. B) Six proteins were significantly increased and four proteins were significantly decreased after candesartan treatment in females.

**Figure 8.**
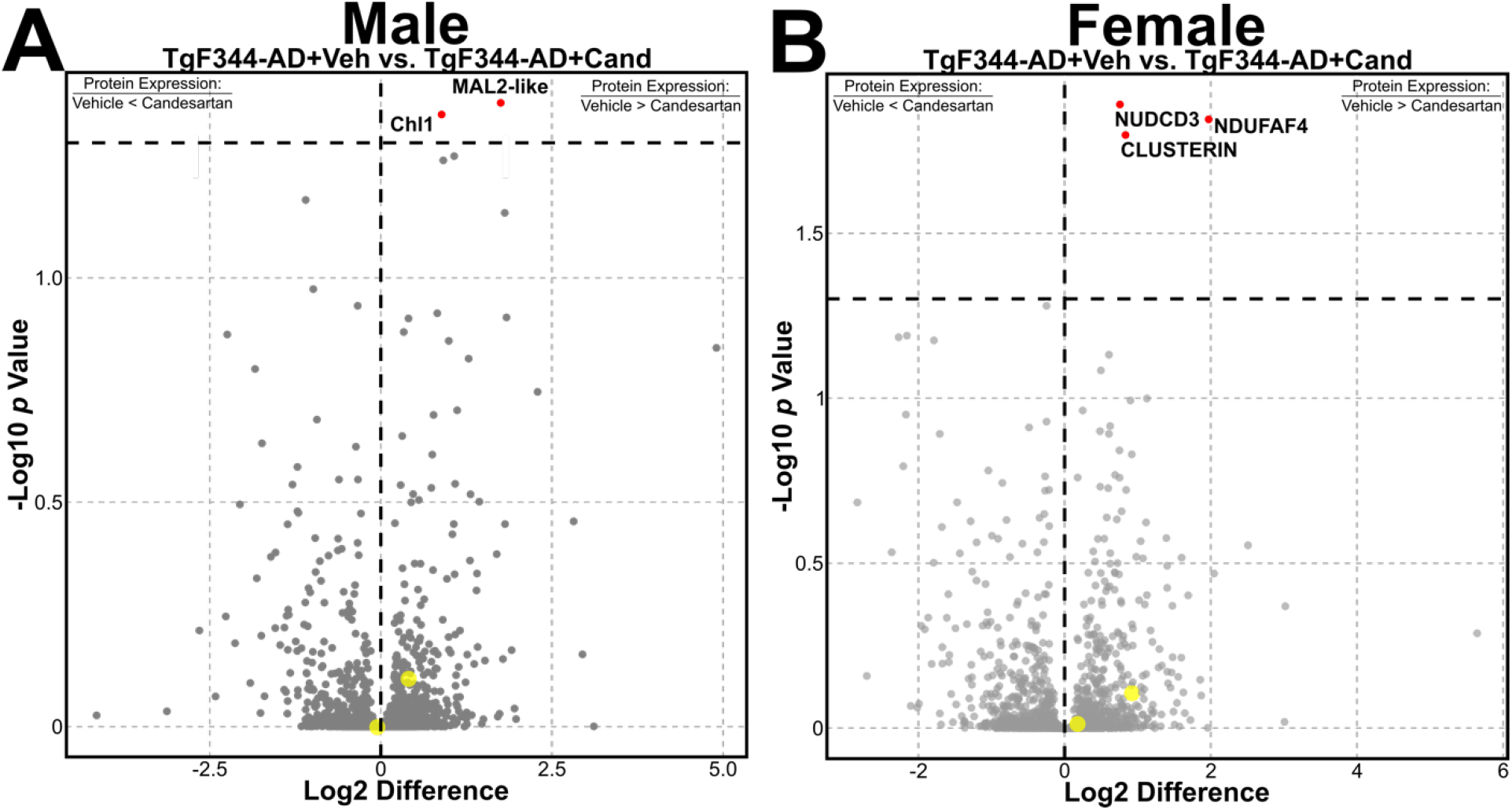
AD-associated proteins were found to have altered expression levels amongst 18-month old WT & TgF344-AD rats. A) There were two proteins that were altered in 18-month old male TgF344-AD rats, Chl1 and MAL2-like protein, but neither protein was found to be significantly altered across all groups via ANOVA (p=0.155 & p=0.154, respectively). B) The expression of three proteins, including the known AD-associated protein clusterin, was changed by treatment with candesartan in 18-month old female TgF344-18 rats. For both volcano plots, yellow dots represent the locations of APP and Midkine.

Untreated TgF344-AD male rats expressed more Chl1 and Mal2-like protein than those treated with candesartan. Three proteins were significantly altered in female TgF344-AD rats treated with candesartan: NDUFAF4, NUDCD3 & clusterin (CLU). CLU has been identified as a late-onset AD-risk gene by independent Genome Wide Association Studies[35, 36]. The difference in relative abundance of each of these proteins were compared between female TgF344-AD+Veh and TgF344-AD+Cand (Figure 7). Pairwise comparison confirmed that there were significant differences in the means between groups for all three proteins [NDUFAF4: *p*=0.014; NUDCD3: *p*=0.013; clusterin: *p*=0.016].

## Discussion

Our results demonstrate that candesartan administered for six months prior to the onset of behavioral manifestations reduces the degree of cognitive impairment in female, but not male, TgF344-AD rats. Candesartan acts as an antagonist of the AT1 receptor, blocking vasoconstriction mediated by angiotensin II and causing vasodilation and reduction in blood pressure. Six months of treatment with candesartan was effective to decrease mean arterial pressure in both wild type and TgF344-AD rats. Candesartan treatment improved performance on the WRAM and decreased the abundance of the AD-risk protein clusterin in female rats only. Women are at a higher risk for AD than men and estrogen has been suggested to be protective against both cardiovascular disease and AD[14, 15]. The risk for major coronary disease and hypertension rises dramatically following menopause[15, 37]. Estradiol appears to downregulate AT1 receptor translation[38] and reduce angiotensin converting enzyme[39], providing a potential RAS-mediated mechanism for the premenopausal protective effect of estrogen on cardiovascular health.

Chronic treatment with candesartan altered clusterin protein expression in female TgF344-AD rats. Clusterin is an extracellular chaperone protein that has been identified as the third greatest genetic risk factor for sporadic AD[40]. The nature of interactions between clusterin and amyloid-β in the pathogenesis is a subject of debate. Under normal conditions, clusterin is thought to provide neuroprotection from amyloid-β by binding to suppress plaque formation. Clusterin is also likely to play a role in amyloid-β clearance through the blood brain barrier, a process mediated binding with LRP2[40]. However, in AD clusterin levels rise in correlation with amyloid-β and appear to contribute to the formation and toxicity of soluble oligomeric amyloid-β peptides[41]. Studies into the interactions between clusterin and the RAS in the brain are lacking, however a number of groups have reported on the relationship between clusterin and angiotensin receptor blockers in the kidney. Clusterin levels have been shown to decrease following the ARBs valsartan[42] and irbesartan[43]. Additionally, amyloid-β has been shown to increase the amount of intracellular clusterin and expression of DKK1, an antagonist of the canonical wnt-signaling pathway[40]. DKK1 signaling through the wnt-PCP-JNK results in the upregulation of genes that promote AD neuropathology and neurotoxicity[44]. Amyloid-β associated upregulation of DKK1 signaling is reliant on clusterin and downregulation in clusterin has been shown to reduce wnt-PCP-JNK signaling and neurotoxicity. Angiotensin II also upregulates DKK1 through AT1R signaling, an effect which is blocked by ARBs[45].

At 12-months, the TgF344-AD rat displayed significant amyloidosis and increases in GFAP levels throughout the brain but appeared cognitively normal on the spontaneous alternation test and novel object recognition test. Multiple investigators have replicated these findings, suggesting that this model develops amyloid pathology with a stereotyped progression beginning at 6-months and continuing through 16-months[21, 46, 47]. At 12- and 18-months, our quantification of amyloid-β supports this progression. We did not demonstrate significant changes in tau protein hyperphosphorylation at 12- or 18-months.

TgF344-AD rats displayed an overall increase in GFAP expression at 12-months and increased GFAP in dorsal hippocampus at 18-months. A natural increase in GFAP levels is expected with advancing age in rats [48], but we found that levels of GFAP in the hippocampus remained consistent from 12- to 18-months in TgF344-AD rats. In AD, GFAP levels typically increase along with increases in amyloid plaques[49, 50]. Astrocyte activity can be an important mediator of amyloid deposition, but chronically activated astrocytes in AD can become neurotoxic[49]. LFQ protein abundance demonstrated a significant increase in GFAP in frontal cortex in TgF344-AD rats. Candesartan treatment appeared to increase GFAP levels in male TgF344-AD rats, while decreasing GFAP in females.

Candesartan did not appear to induce differential effects in vascular function between TgF344-AD and WT rats. While we cannot state definitively that AngII effects on the vasculature in the brain are not contributing to the decline in cognitive behaviors, it is clear that there are no significant differences between the WT and TgF344-AD in baseline blood pressure and vascular function and if anything, the TgF344-AD model demonstrated improved vascular function at both 12- and 18-months. Furthermore, candesartan treatment similarly impacted the WT and TgF344 rats with respect to blood pressure and vascular function. The effects of six months of candesartan treatment on vascular function suggest that chronic exposure to candesartan can affect sensitivity to vasoconstrictors and vasodilators regardless of genotype or sex of the rat. Further studies are needed to define the mechanisms underlying the long-term candesartan-specific effects on vascular function which may be related to factors such as, improved mitochondria function resulting from AT1 receptor blockade [51].

At 18-months, both male and female WT rats reduced their latency to complete the maze and the total number of maze errors over the course of training. Vehicle-treated male and female TgF344-AD rats displayed learning impairments on the WRAM at 18-months. While male and female TgF344-AD rats were able to complete the maze in less time with training, neither group decreased their cumulative maze errors on the WRAM over 12 days of testing. After candesartan administration, female TgF344-AD were able to decrease their cumulative maze errors on the final training sessions. Improvement in maze performance following candesartan treatment coincided with a reduction in the relative abundance of clusterin and GFAP in frontal cortex. We did not demonstrate working or reference memory deficits in TgF344-AD rats at 12-months of age. While some studies have shown deficits in reversal learning as early as six months and impaired performance on learning and reference memory tasks at 10- to 12-months of age when compared to WT rats[52-54], other groups have found no difference between WT and TgF344-AD rats on learning and memory tasks at or beyond 12-months of age[55-57].

This study demonstrates an effect of the angiotensin II receptor blocker, candesartan, to improve learning and spatial memory in aged female TgF344-AD rats. The advantages of this approach are the large sample cohort and the assessment of the brain proteome in TgF344-AD rats as well as age-matched and treatment-matched control littermates. Additionally, this study is unique in its assessment of long-term administration of an angiotensin receptor blocker on vascular contractility. Six months of candesartan treatment was well tolerated in both WT rats and the TgF344-AD rats, and led to similar effects on vascular function in both models. WT and TgF344-AD rats entered the study at the age of 12-months. While these rats were not displaying behavioral differences from WT littermates at this age, TgF344-AD rats had already developed a significant amount of amyloid-β deposition and an increase in GFAP. Candesartan did not appear to slow the aggregation of amyloid-β when administered between 12- and 18-months. While we did see a sex-dependent effect of candesartan on GFAP levels at 18-months in TgF344-AD rats, the size of these effects are small and difficult to interpret, especially given the consistent increase in GFAP from 12-months. We cannot, from our data, draw conclusions about candesartan’s potential to reduce amyloid burden and further influence GFAP levels if ARB administration starts before significant aggregation. Additionally, the female rats in our study were ovary-intact throughout their lifespan. The majority of rats that remain ovary intact into adulthood maintain high levels of 17β-estradiol throughout estropause[58]. Additionally, advanced age has not been shown to modulate the expression of AT1 or AT2 receptors in rats[59]. While we did not track estrus for our female rats, it is unlikely that the effect of candesartan is dependent on reduction in estradiol levels in our rats. The sex difference in response to candesartan is interesting to note given the interactions between estradiol and the RAS. However, there are notable differences between menopause in humans and estropause in rodents. The results of this study demonstrate the potential for angiotensin receptor blockers to delay the presentation of the behavioral symptoms of AD when administered in the early disease phase. Further studies are needed to determine the biological mechanisms underlying the effects of candesartan on AD disease progression.

